# Connexin 43 mediated monocyte mitochondrial transfer prevents cisplatin induced sensory neurodegeneration

**DOI:** 10.1101/2024.10.24.620120

**Authors:** Rebecca Owen, Jack Corbett, Mark Paul-Clark, Richard Philip Hulse

**Affiliations:** Centre for Systems Health and integrated Metabolic Research, Department of Biosciences, School of Science and Technology, Nottingham Trent University, Nottingham, NG11 8NS

**Keywords:** dorsal root ganglia, mitochondria, chemotherapy, cisplatin, neurodegeneration, inflammation, macrophage, neuropathy, mitochondria, gap junction, connexin 43

## Abstract

Platinum based chemotherapeutics including cisplatin are front-line treatments for paediatric and adult cancer. Despite advancements in medical interventions, chemotherapy-induced peripheral sensory neuropathy is a common adverse health related complication that can persist for the long-term and impacts upon an individual’s quality of life. Recently, the causes of chemotherapy induced sensory neurodegeneration has been linked to sensory neuronal mitochondrial dysfunction. Here this study investigated how monocytic mitochondria donation to recipient cisplatin damaged dorsal root ganglia (DRG) sensory neurons prevented platinum-based chemotherapy-induced sensory neurotoxicity. Neuronal cell line, SH-SY5Y, or mouse DRG sensory neurons were treated with either vehicle or cisplatin, and co-cultured with mitotracker-labelled THP1 monocytes. Cisplatin induced dysmorphic mitochondria and diminished oxidative phosphorylation dependent energy production in cisplatin treated dorsal root ganglia sensory neurons. DRG sensory neurons exposed to cisplatin were recipients of monocyte mitochondria indicated by increased intracellular mitotracker fluorescent labelling. Mitochondrial transfer to sensory neurons was neuroprotective, preventing neurite loss and sensory neuronal apoptosis. Vehicle treated DRG sensory neurons did not demonstrate significant mitochondrial uptake. Furthermore, cisplatin induced mitochondrial transfer was prevented by pharmacological inhibition of gap junction protein, connexin 43. Connexin 43 inhibition led to reduced neuroprotective capacity via mitochondrial transfer. These findings demonstrate that monocytic mitochondria transfer to DRG sensory neurons damaged by cisplatin, is dependent upon gap junction intercellular communication to promote sensory neuronal survival. This novel process in sensory neuronal protection is a potential novel therapeutic intervention for alleviating neuropathic pain in individuals treated for cancer.

## Introduction

Cisplatin is a front-line platinum-based chemotherapeutic used for the treatment of adult and paediatric cancer. Recent improvements in chemotherapy regimens have resulted in an increase in 5 year-survival rate from paediatric malignancies to 86% in 2018 [15]. However, chemotherapy also impacts non-cancerous host cells, such as immune cells and sensory neurons. This causes adverse health related complications [3], with cisplatin disrupting the development of physiological tissues and processes leading to longer term health implications in those treated for childhood cancer. Symptoms of CIPN can manifest years after the termination of chemotherapy[22]. This is demonstrated by CIPN affecting 70% of childhood cancer survivors[6] with this increasingly presented with increased patient age. Furthermore, the number of patients affected are unfortunately set to increase with survival rates of paediatric cancers improving. As a consequence, large numbers of adult survivors of paediatric cancers commonly suffer from chemotherapy-induced peripheral sensory neuropathy (CIPN)[4] as well as adult cancer patients. This CIPN commonly manifests as dysesthesia and paresthesia in the extremities[19] that can persist for the long-term[21], with limited effective analgesia available.

Cisplatin induced neuropathic pain is accompanied by sensory neural architectural changes, includes sensory axonal atrophy [1] and intraepidermal nerve fibre remodelling [13], that arise due to sensory neurotoxicity. This sensory neuropathology impedes normal sensory neuronal function, presenting as reduced sensory nerve fibre conduction velocity [16] and nociceptor sensitization [11,27]. As to causation, cisplatin induced alterations in mitochondrial processes have been identified in platinum-based chemotherapy induced neuropathologies[12,14]. Cisplatin forms adducts with mtDNA impairing mitochondrial function and induces morphological abnormalities [18,24]. Dorsal root ganglia sensory neurons place high energy demands upon their own cellular respiration processes compared to other cell types, with alterations in oxidative phosphorylation driving persistent pain states[29]. Therefore sensory neurons are dependent upon efficient functionally active mitochondria to provide the essential energy provision to support action potential generation and energy mobility in the axonal processes to respond to neuronal stress [20]. Consequently, sensory neuronal mitochondrial dysfunction [2] is heavily implicated in the underlying mechanisms of CIPN and nociceptor sensitization [5], as well as due to the increased susceptibility to oxidative stress [10,16].

This study will investigate if the uptake of monocyte-derived mitochondria recovers neuronal architecture and function. Finally, this study will elucidate new neuro-modulatory pathways involved in the transfer of mitochondria from monocytes to cisplatin-damaged sensory neurons, in order to identify a novel therapeutic target for alleviating chronic neuropathic pain in adult survivors of paediatric chemotherapy treatment.

## Methods

### Ethical approval and animals used

All studies involving rodents were performed under the Animal (Scientific Procedures) Act 1986 and authorisation in line with UK Home Office licenses and ARRIVE guidelines. Studies were approved following consultation with local Animal Welfare and Ethics Review Board at Nottingham Trent University. Animals had ad libitum access to standard chow and were housed under 12:12h light:dark conditions.

### Immortalised Cell culture

Neuroblastoma cell line SH-SY5Y were maintained in Dulbecco’s Modified Eagle Medium (DMEM), supplemented with Fetal Bovine Serum (FBS) (10%), Penicillin and Streptomycin (Pen-Strep) (1%), and Sodium Pyruvate (1%. Cells were cultured until confluent and subsequently plated in a 24-well plate at a cell density of 5,000 cells per well. Monocyte THP-1 cells were maintained in Rodwell Park Memorial Institute (RMPI) media, supplemented with FBS (10%) and Pen-Strep (1%).

### Primary DRG Sensory Neuronal Cell Culture

Mouse (5 C57.Bl6J postnatal ∼day 7 mice used per assay) Dorsal Root Ganglia (DRG) neurons were maintained in Ham’s F12 (Gibco) medium, supplemented with N2 (1x, Gibco 17502-048), Bovine Serum Albumin (BSA 1%, Sigma A9576), and Pen-Strep (2%). Using either a 24-well plate or 8 well imaging chambers (Ibidi), wells were coated in Poly-L-Lysine (Sigma P4707) overnight. Poly L-lysine was removed, and Laminin (Sigma L2020, 1µg/ml) applied for 2 hours. Isolated DRG neurons were enzymatically dissociated with collagenase type XI (Sigma C9407, 0.0125% for 2 hours. DRG neurons mechanically dissociated and centrifuged across two 15% BSA cushions, at 1000 relative centrifugal force (rcf) for 16 minutes. Cell pellets were resuspended in media, counted and plated in either a 24-well plate or glass chambers at 7,000, and 5,000 DRG neurons per well, respectively. DRG neurons were allowed to grow for one week.

### Drug Preparation and Experimental Treatments

Either DRG neurons or SH-SY5Y cells were treated with either vehicle or cisplatin (5µg/ml, Tocris 2251 5A/256231) for either 2 or 24 hours. THP-1 cells labelled with Mitotracker Red CMXRos (1:10,000) and were added to either DRG sensory neurons or SH-SY5Y cultures at 5,000 per well. In some instances, THP-1 cells treated with cisplatin (5µg/ml) for 2 hours were added to DRG cultures. SH-SY5Y and DRG sensory neurons were treated with increasing concentrations of TAT Gap19 (Tocris 2267 9A) (0, 0.1, 1, 7, and 10µM) were pretreated for either 2 or 24 hours. SH-SY5Y cells were also labelled with Mitotracker Red CMXRos (1:10,000) in some instances.

### Immunohistochemistry

DRG neurons were cultured on glass coverslips in a 24 well plate and experimental treatments were applied. Cells were subsequently washed with Phosphate Buffered Saline (PBS) solution for 5 minutes and fixed using Paraformaldehyde (1%) for 15 minutes at room temperature. DRG neurons were washed with PBS (twice for 5 minutes), and then permeabilised using PBS 0.2% Tritonx-100 for 5 minutes Coverslips were incubated in blocking solution (PBS 0.2% Triton x-100, 5% BSA) for 30 minutes at room temperature. DRG neurons were incubated at 4oC overnight with rabbit (Abcam Ab18207) or guinea pig (SySy) β3 tubulin (1 in 500), in some instances Phalloidin (AlexaFlor®488 A12379) (1µM) or rabbit anti connexin 43 (Cell signalling). Coverslips were washed twice for PBS for 5 minutes. Coverslips were incubated with Alexafluor 488 or 647 donkey anti-rabbit IgG (A31571) (1 in 1000) or Alexafluor 555 donkey anti-guinea pig IgG for 2 hours at room temperature. Coverslips were washed twice with PBS for 5 minutes, and incubated with DAPI (Abcam Ab228549) (1 in 1000) for 5 minutes. Each coverslip was mounted onto glass slides using Vectashield Antifade Mounting Medium. The LEICA confocal microscope was used to visualise DRG neurons.

### Live cell imaging

The LEICA Thunder microscope was used to visualise Mitotracker Red CMXRos and Brightfield in either SH-SY5Y or DRG neurons. Images of brightfield and Mitotracker fluorescence (555) were acquired over 30 minutes, 1 hour, or 15 hours. Either magnification X20 or X63 oil immersion was used. 3 regions of interest were acquired per well of either a 24-well plate or removable glass slides.

### Gene expression analysis

Cisplatin-induced neuropathic pain was established in neonatal Wistar rats that were administered via intraperitoneal injection either vehicle (phosphate-buffered saline; PBS) or cisplatin (Sigma-Aldrich; 0.3 mg/kg) on postnatal days 14 and 16. Rodents were weaned postnatal day ∼21 and housed according to sex. Lumbar 2-5 DRGs were identified and extracted at P23 and immediately frozen. RNA sequencing was performed by Novogene as previously outlined[11], and data was processed from available from data repository GSE281897. Differential gene expression analysis was performed on raw counts using the R statistical package DESeq2 (v1.20.0). Pathway analysis was performed in this studies exploring genes with a >1.5-fold increase in expression [fold change (FC)≥1.5] and P<0.05 were considered significantly upregulated, and genes with a >1.5-fold decrease in expression (FC≤1.5) and P<0.05 were considered significantly downregulated. Non-coding genes were filtered out of the analyses. Gene enrichment pathway analysis was performed using STRING with the most abundant biological processes and molecular function highlighted, and a false discovery rate of <0.05 applied to adjusted P-values.

### Statistical Analysis

Analysis was carried out using image J (Fiji), Microsoft Excel, Imaris software, and GraphPad Prism 10 software. All data are presented as mean ± SEM unless otherwise stated. Mitotracker functionality; Image J or Imaris was used to quantify Mitotracker functionality in either SH-SY5Y or DRG neurons. 3 images were acquired per well of either a 24-well plate or removable glass chambers, and 3 regions of interest were analysed per image. The integrated density of Mitotracker fluorescence was measured in either SH-SY5Y or DRG neurons by using the freehand tool to draw around the circumference of either SH-SY5Y or DRG neurons in the Mitotracker fluorescence channel. Additionally, mitochondria segmental length and volume were determined using Imaris. Results were normalized to that of the vehicle for each experiment. Neurite outgrowth; Neurite outgrowth was measured using Image J. Parameters for neuronal identification were determined by the presence of β3 tubulin staining, and neurite outgrowth was determined as the length of phalloidin staining from the cell body. This was achieved by using the freehand line draw tool to draw from the phalloidin staining of the DRG cell body to the end of the neurite. A two-way ANOVA and a Dunnett’s post hoc test was used to analyse change in Mitotracker fluorescence. Either a one or two way ANOVA was used to determine significance in neurite outgrowth in DRG neurons co-cultured with Thp-1 following cisplatin treatment compared to vehicle. P values are represented as *P<0.05, **P<0.01, ***P<0.001, ****P<0.0001, and NS not significant.

## Results

Rodents administered cisplatin early in life demonstrate disturbances in DRG sensory neuronal mitochondrial functionality and neuritogenesis in adulthood (Fig.1A&B). Adult lumbar dorsal root ganglia sensory neurons that were exposed to cisplatin during infancy had a diminishment in aerobic ATP generation and oxidative phosphorylation capability (Fig.1B). Furthermore, in isolated mouse DRG sensory neurons cultured in vitro mitochondria were visualized (Fig.1C&D), and following cisplatin treatment there was a reduction in mitochondrial size; mitochondrial fragmented volume (Fig.1E) and length (Fig.1F). Similarly, in the neuroblastoma cell line, SH-SY5Y, following treatment with cisplatin, there was a reduction in mitotracker fluorescence when compared to vehicle (Fig. 2B).

**Figure 1:**
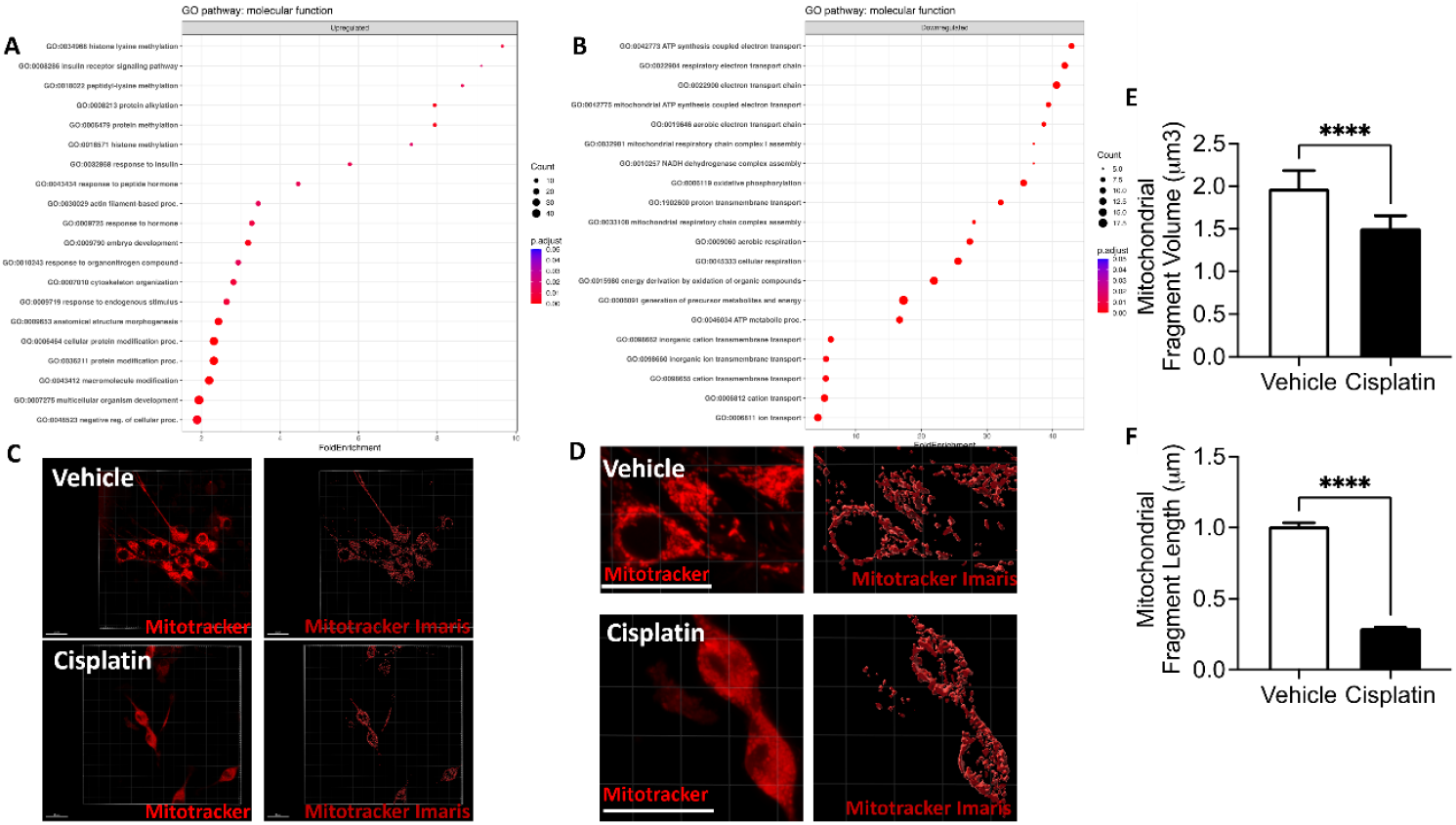
Cisplatin induced mitochondrial disturbances in dorsal root ganglia sensory neurons. Wister-Hans rats were treated with 0. 3mg/kg cisplatin or saline (control) on postnatal days 14 and 16. DRGs were extracted 7 days later, with gene ontology analysis for molecular function of [A] upregulated and [B] downregulated differentially expressed genes between vehicle and cisplatin treated rodents. Cisplatin induced changes in DRG sensory neuronal mitochondrial morphology. Representative images of DRG sensory neurons [C low power; D high power] treated with either vehicle or cisplatin (5µg/ml) and fluorescently labelled with Mitotracker CMXRos (Scale bar = 20µm). Mitochondrial morphology was investigated using Imaris software demonstrating decreased neuronal mitochondrial fragmented [E] volume and [F] length was observed in cisplatin treated DRG sensory neurons compared to vehicle. Unpaired-t-test, *** P<0.001, **** P<0.0001.

**Figure 2:**
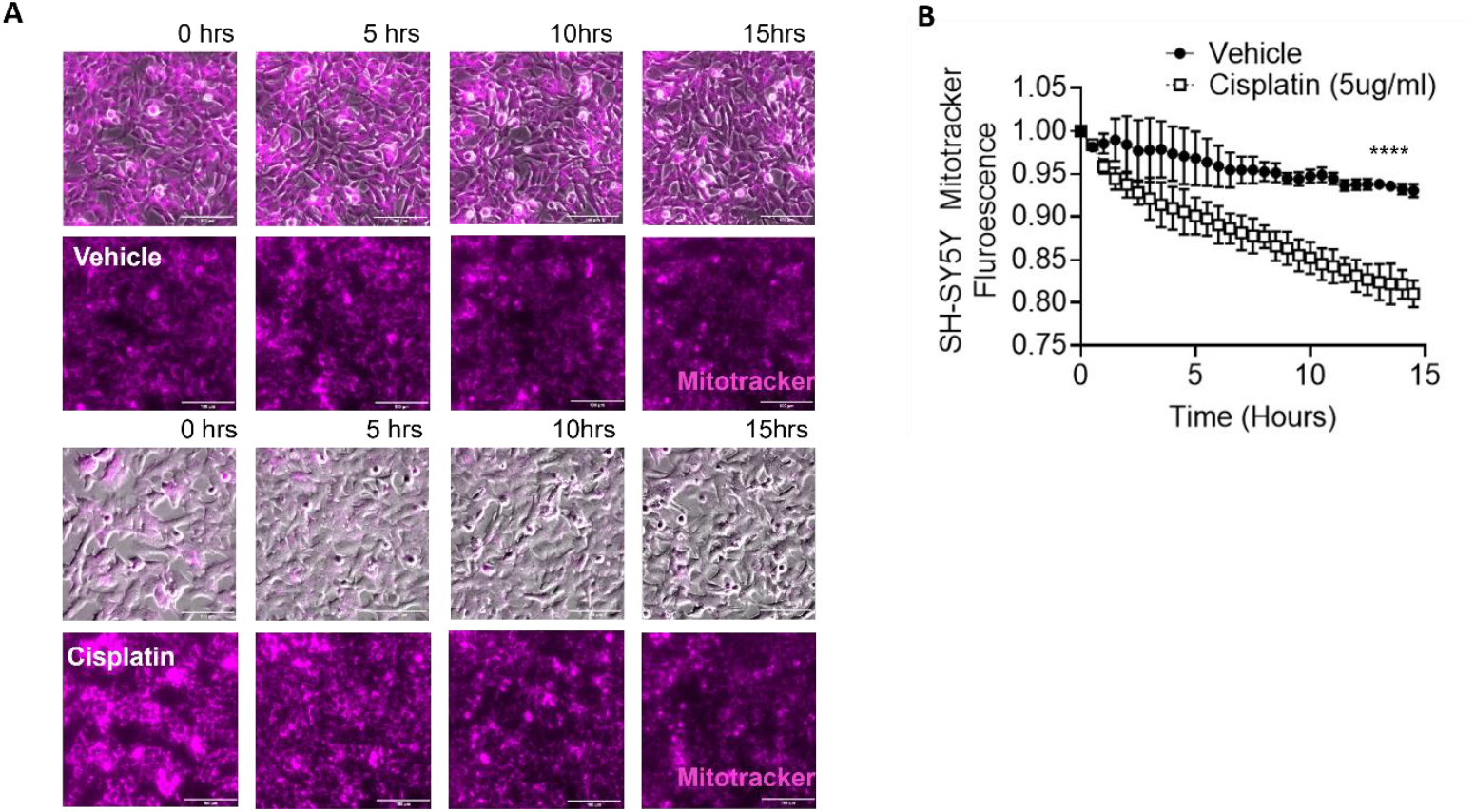
Cisplatin causes mitochondrial dysfunction in SH-SY5Y cells. [A] SH-SY5Y cells were either treated with either vehicle or cisplatin (5µg/ml) for 2 hours, and mitotracker functionality was measured for 15 hours (Scale bar = 100µm). [B] A decrease in mitotracker fluorescence was observed in cisplatin-treated SH-SY5Y cells compared to vehicle over time. Two-way ANOVA and a Kruskal Wallis test for multiple comparisons, **** p<0.0001. Error bars are mean±SD.

Recently, preservation (REF) or rescue (REF) of DRG sensory neuronal mitochondrial preservation has been shown to be integral in modulating nociceptive processing. Monocytes transfer mitochondria to other cell types (REF) and here we investigated whether this occurred in DRG sensory neurons. SH-SY5Y cells were treated with either vehicle or cisplatin and co-cultured with THP1 monocytic cells, which were also either treated with vehicle or cisplatin (Fig.3A). An increase in mitotracker fluorescence was observed in SH-SY5Y cells that had been treated with cisplatin, when co-cultured with vehicle treated monocytes (Fig.3B). This mitochondrial transfer from monocytes to neurons occurred within 20 minutes (Fig.3C). However, there was no indication of mitochondrial transfer from monocytes to vehicle treated DRG sensory neurons (Fig.3C). Similarly, the transfer of mitochondria from monocytes to cisplatin treated DRG sensory neurons was also demonstrated (Fig.4A). Mitotracker labelled THP1 cells were co-cultured with DRG sensory neurons and an increase in fluorescence was observed in DRG sensory neurons treated with cisplatin (Fig.4B). There was no change in monocyte mitochondrial transfer in vehicle treated DRG sensory neurons (Fig.4B). Additionally, cisplatin induced sensory neurodegeneration indicated by reductions in neurite length (Fig.5A), neurite number (Fig.5B) and total neuron growth increasing (Fig.5C), which was prevented by coculturing with THP1 monocytes (Fig.5D). Furthermore, cisplatin treatment led to a reduction in OCR values in DRG sensory neurons, which was rescued by coculturing with THP1 monocytes (Fig5.E).

**Figure 3:**
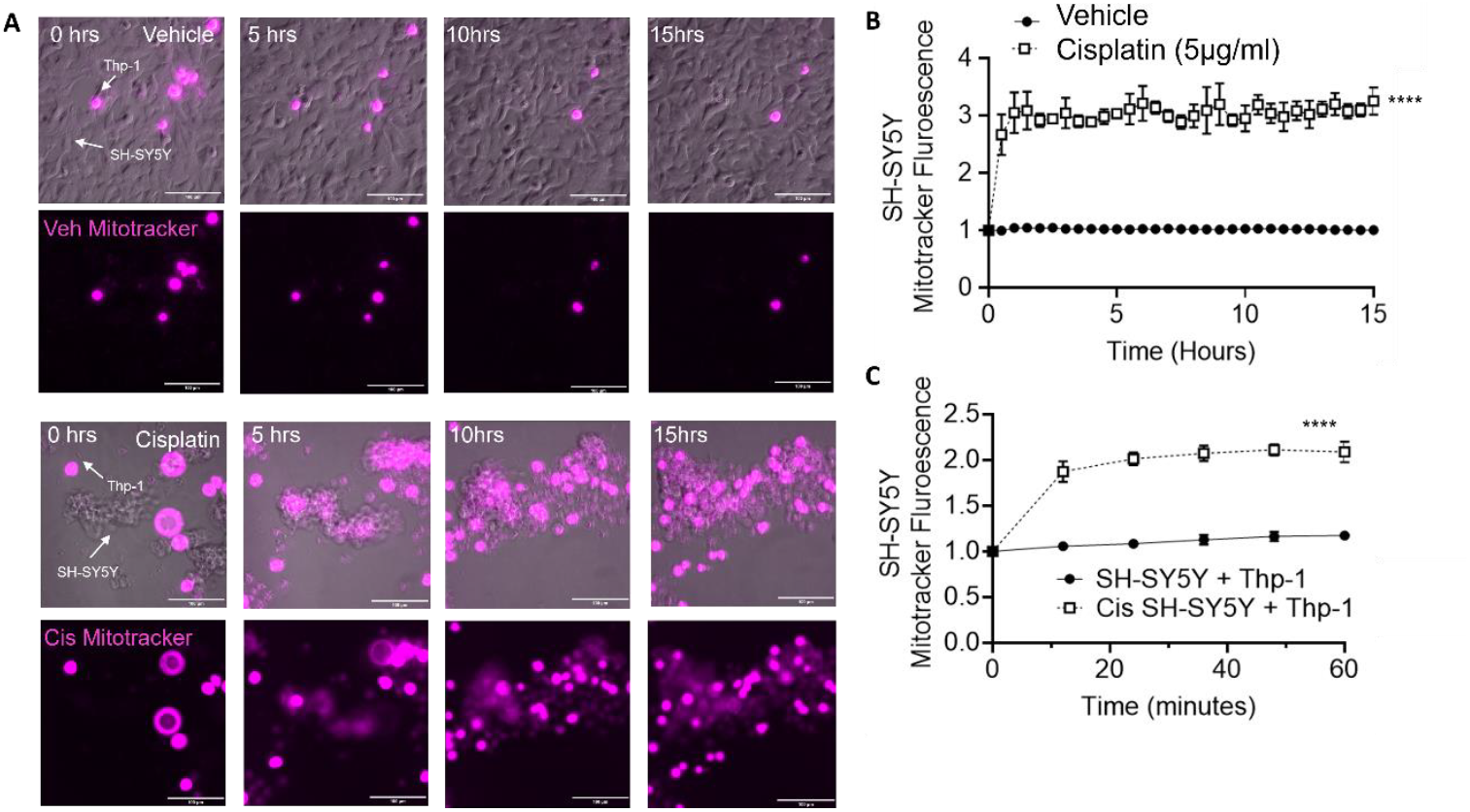
Monocytes transfer mitochondria to cisplatin-treated neurons. [A] SH-SY5Y cells were treated with either vehicle or cisplatin (5µg/ml) for 24 hours and co-cultured with THP-1 cells fluorescently that were labelled with mitotracker (Scale bar; 100µm). [B] When co-cultured with THP-1 cells, there was an increased uptake of mitotracker labelled mitochondria in cisplatin-treated SH-SY5Y cells compared to vehicle, whereas there were no difference in vehicle treated SH-SY5Y cells. [C] There was an increased uptake of mitotracker-labelled mitochondria in neuronal like cells within 20 minutes when vehicle-treated THP-1 cells were co-cultured with cisplatin treated SH-SY5Y cells. There was no difference in vehicle treated SH-SY5Y cells. **** p<0.0001, Two-way ANOVA and a Kruskal Wallis test for multiple comparisons. Error bars are mean±SD.

**Figure 4:**
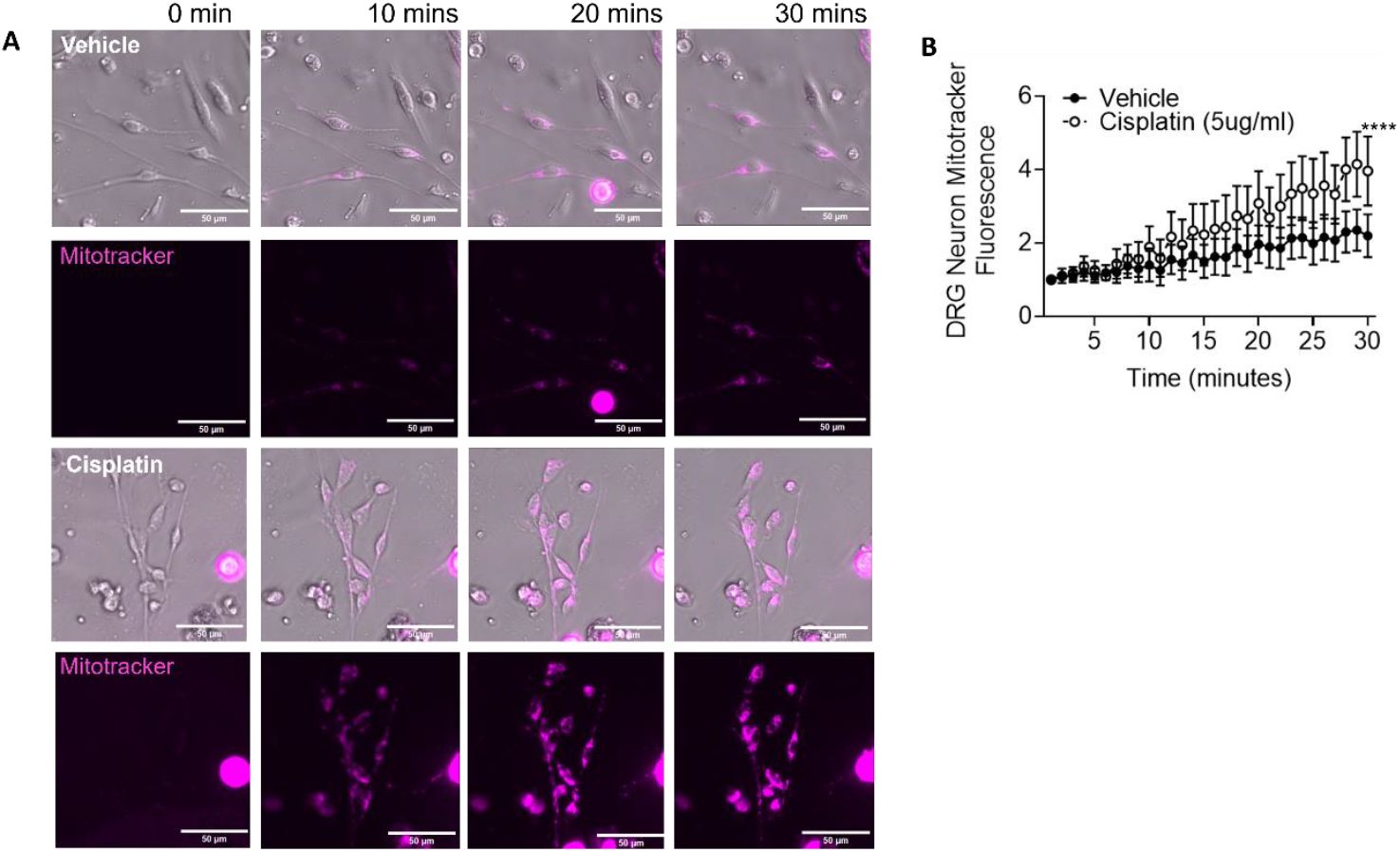
Monocytes transfer mitochondria to cisplatin treated DRG sensory neurons. [A] Isolated Dorsal Root Ganglia (DRG) sensory neurons (Neonatal P7 mice) were treated with either vehicle or cisplatin (5µg/ml) and co-cultured with THP-1 cells fluorescently labelled with mitotracker. [B] When co-cultured with THP-1 cells, there was an increased uptake of mitotracker labelled mitochondria in cisplatin-treated DRG neurons compared to vehicle. **** p<0.0001 at t=12 mins, two-way ANOVA and a Kruskal Wallis test for multiple comparisons. Scale bar; 50µm. Error bars are mean±SD.

**Figure 5:**
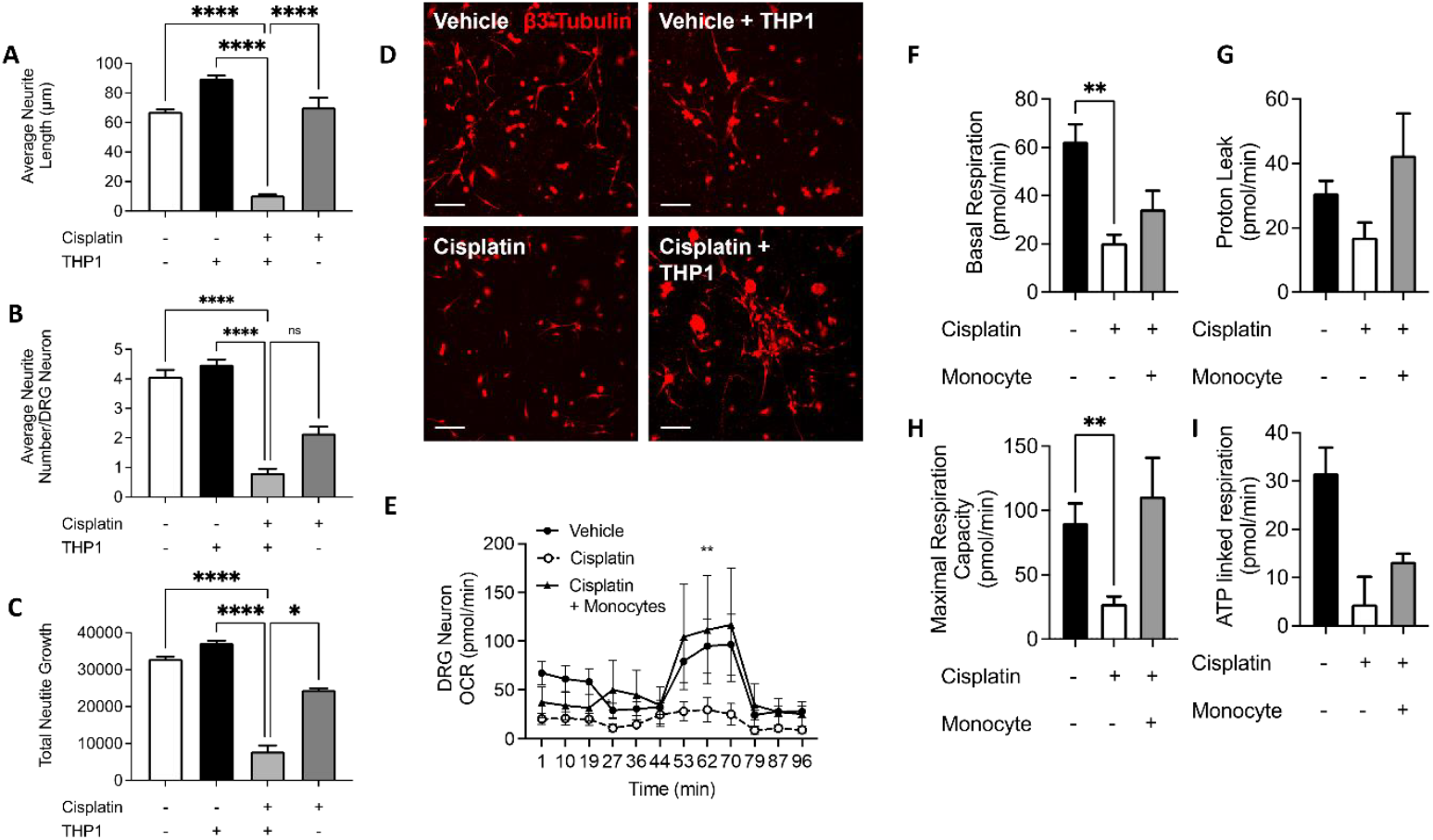
Mitochondrial transfer from monocytes to DRG sensory neurons prevents cisplatin induced neurodegeneration and rescues mitochondrial respiration. Isolated DRG sensory neurons were treated with either vehicle or cisplatin (5µg/ml) and co-cultured with THP-1 cells overnight. [A] Average neurite length, [B] number and [C] total neurite growth per neuron was determined, with [D] monocyte mediated mitochondrial transfer preventing cisplatin induced sensory neurodegeneration (Red β3 tubulin, scale bar = 100µm). [E] Oxygen consumption rate was determined in DRG sensory neurons with and without cisplatin plus co-culturing with THP1 monocytes to explore basal respiration, [F] proton leak, [G] maximal respiration [H] capacity and [I] ATP linked respiration. Donation of monocyte mitochondria prevented cisplatin induced reduction in basal and maximal respiration capacity. *P<0.05, **P<0.01, *** P<0.001, **** p<0.0001 one-way ANOVA and a Kruskal Wallis test for multiple comparisons.

The process that mitochondria transfer between cells has been reported by varying pathways, with connexins a mediator of intercellular organelle transfer[8] (REF). In DRG sensory neurons gap junction protein connexin 43 (CX43) was identified (Fig.6A). Following induction of cisplatin induced sensory neurodegeneration (Fig.6B), there was an associated increase in expression of connexin 43 in sensory neurons that was associated with increased sensory neurodegeneration (Fig.6C&D). Consequently, pharmacological connexin 43 inhibition was adopted to investigate the effects on the transfer of mitochondria from monocytes to cisplatin damaged sensory neurons. SH-SY5Y were treated with vehicle or cisplatin in combination with mitotracker labelled THP1 monocytes and in the presence of TAT Gap-19, connexin 43 inhibitor (Fig.7A). A dose-dependent decrease in mitotracker fluorescence was observed in cisplatin treated SH-SY5Y cells following Gap-19 treatment (Fig.7B), whereas there was no difference in vehicle treated SH-SY5Y cells. In DRG sensory neurons (Fig. 8A) the transfer of mitochondria from monocytes to cisplatin damaged sensory neurons results was dependent upon connexin 43, with Gap-19 treatment inducing a dose-dependent decrease in mitotracker fluorescence in cisplatin treated DRG sensory neurons (Fig. 8B). GAP 19 treatment did not cause any change in There was no mitochondrial transfer in vehicle treatment DRG sensory neurons, which was unaltered by connexin 43 inhibition (Fig. 8C). Subsequently it was identified that connexin 43 mediated monocyte mitochondrial donation to cisplatin damaged DRG sensory neurons prevented sensory neurodegeneration (Fig.9A). There was an increase in neurite outgrowth in cisplatin treated DRG sensory neurons following monocyte mitochondria donation, which dose dependently reduced with increasing concentrations of TATGap19 (Fig.9B). In addition, monocyte donated mitochondria prevented cisplatin induced DRG sensory neuronal cell death, which was prevented by connexin 43 inhibition (Fig.9C).

**Figure 6:**
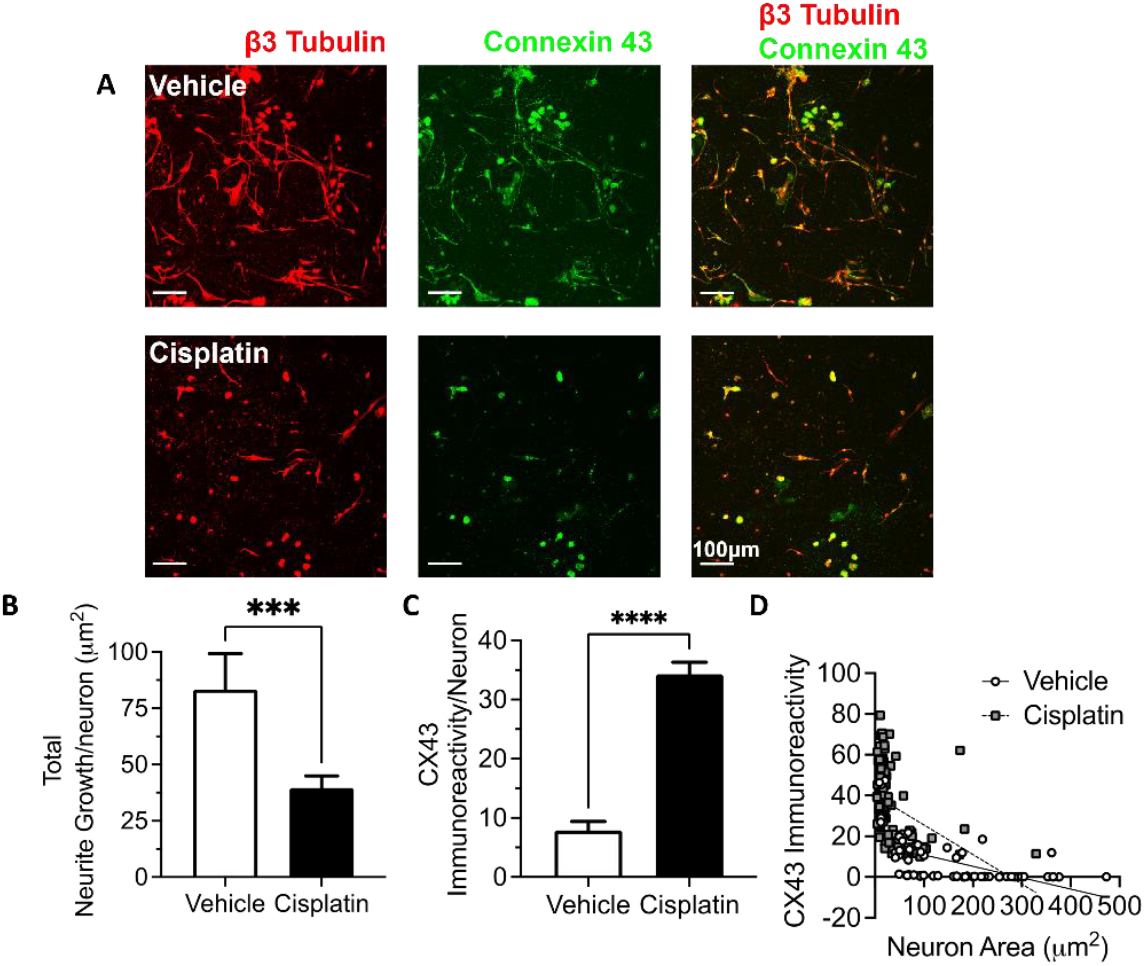
Connexin 43 expression in DRG sensory neurons is associated with cisplatin induced sensory neurodegeneration. [A] Representative images (Scale bar = 100µm) of isolated mouse DRG sensory neurons treated with either vehicle or cisplatin, demonstrating expression of connexin 43. [B] Cisplatin induced sensory neurodegeneration demonstrated by reduced neurite growth. [C] Connexin 43 was increased following cisplatin treatment, [D] with increased expression identified on DRG sensory neurons that had less neurite growth. *** P<0.001, **** p<0.0001 one-way ANOVA and a Kruskal Wallis test for multiple comparisons.

**Figure 7:**
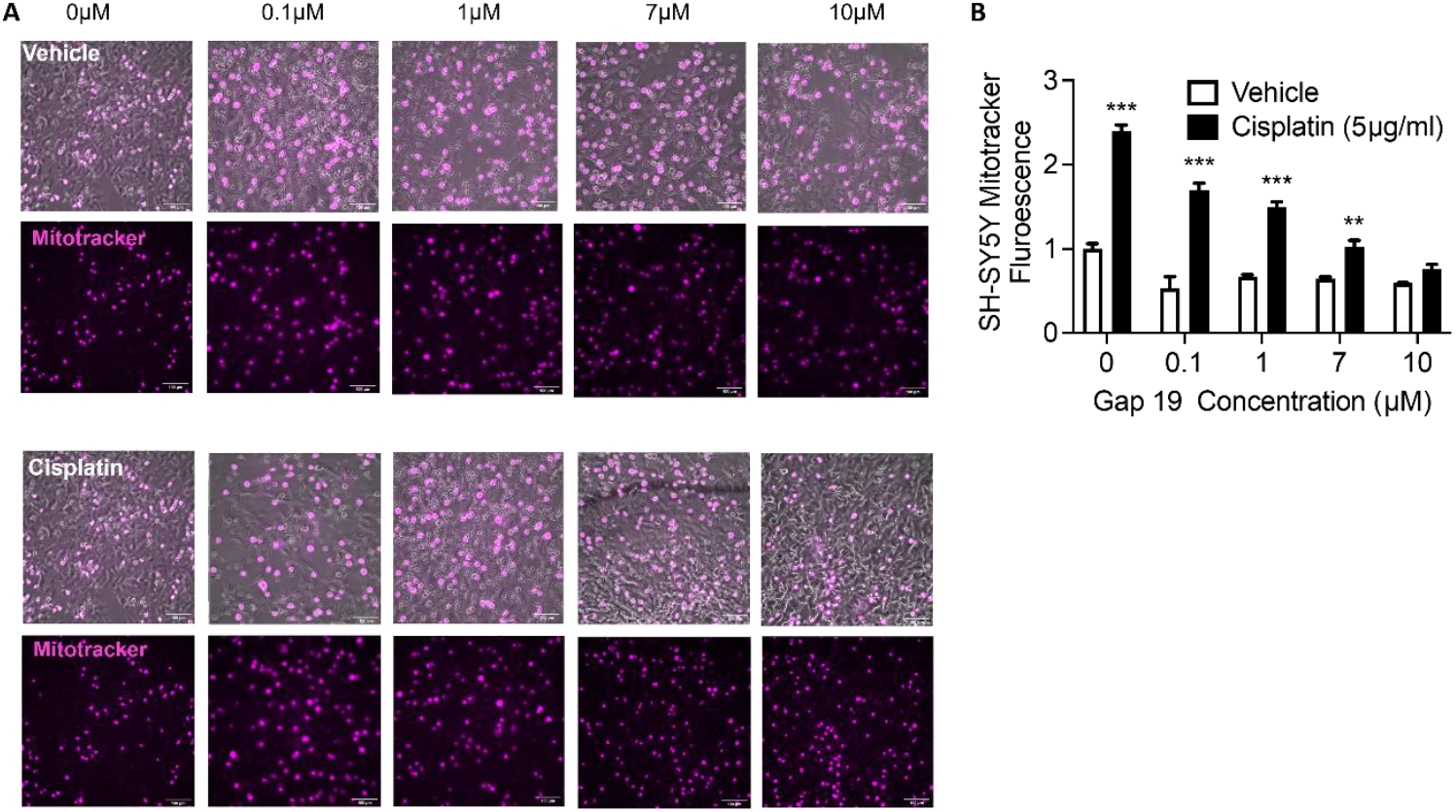
Mitochondrial transfer from monocytes to SH-SY5Y is mediated by connexin 43. [A] SH-SY5Y were treated with either vehicle or cisplatin with increasing concentrations of TATGap-19 and subsequently co-cultured with mitotrackcer labelled monocyte Thp-1 cells (representative images. scale bar = 100µm). [B] TATGap-19 treatment resulted in a decreased uptake of mitotracker labelled mitochondria by cisplatin-treated SH-SY5Y cells. Two-way ANOVA and a Kruskal Wallis test for multiple comparisons. **P<0.01, *** P<0.001. EC50; Veh = 3.582µM, Cis = 7.185µM.

**Figure 8:**
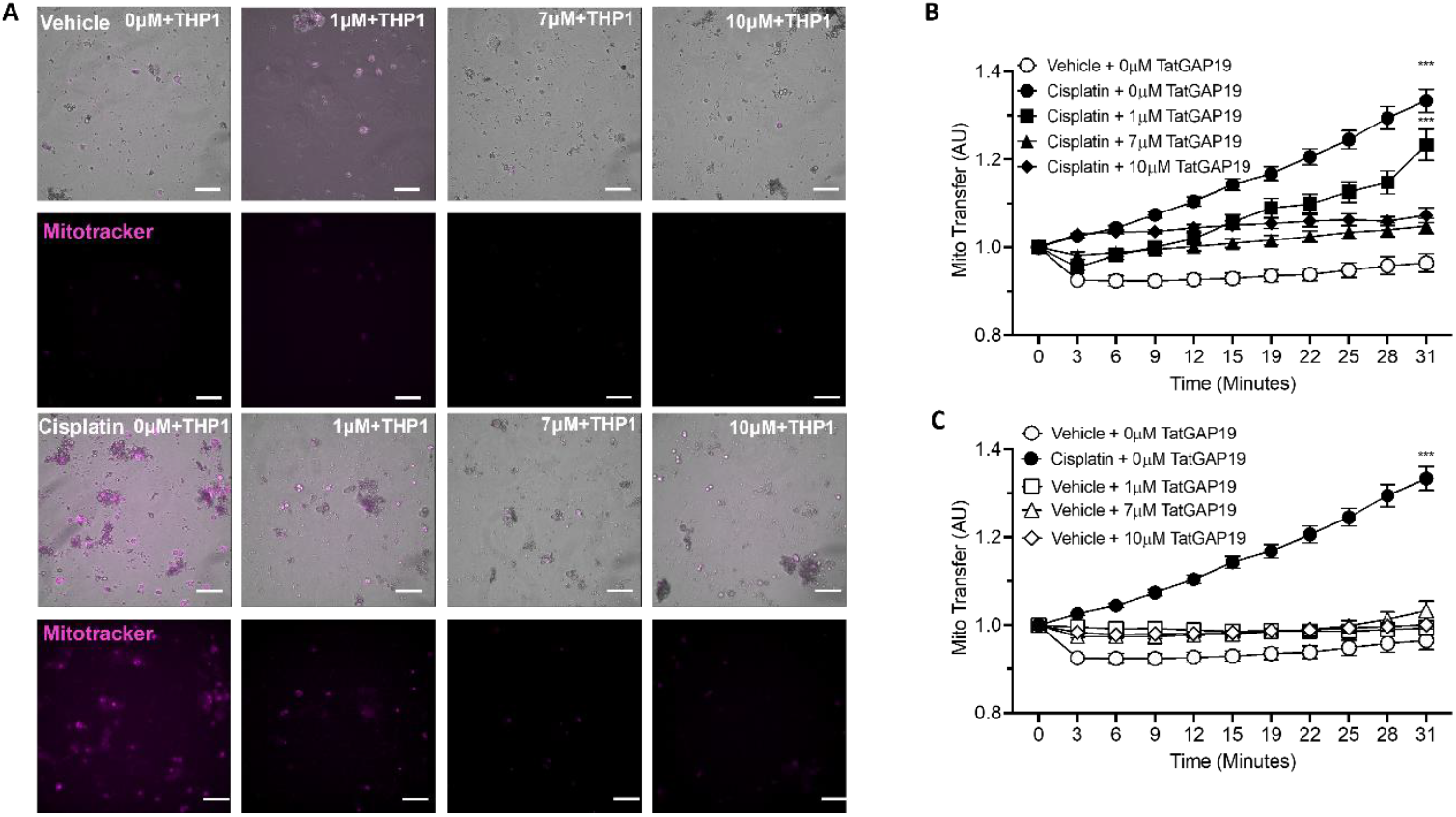
Mitochondrial transfer from monocytes to DRG sensory neurons is mediated by Connexin 43 inhibition. [A] DRG sensory neurons were treated with either vehicle or cisplatin and increasing concentrations of TATGap-19. Subsequently these were co-cultured with mitotracker labelled monocyte THP-1 cells (representative images. scale bar = 100µm). [B] TATGap-19 treatment resulted in a decreased uptake of mitotracker labelled mitochondria by cisplatin treated DRG sensory neurons. Two-way ANOVA and a Kruskal Wallis test for multiple comparisons. *** p<0.001.

**Figure 9:**
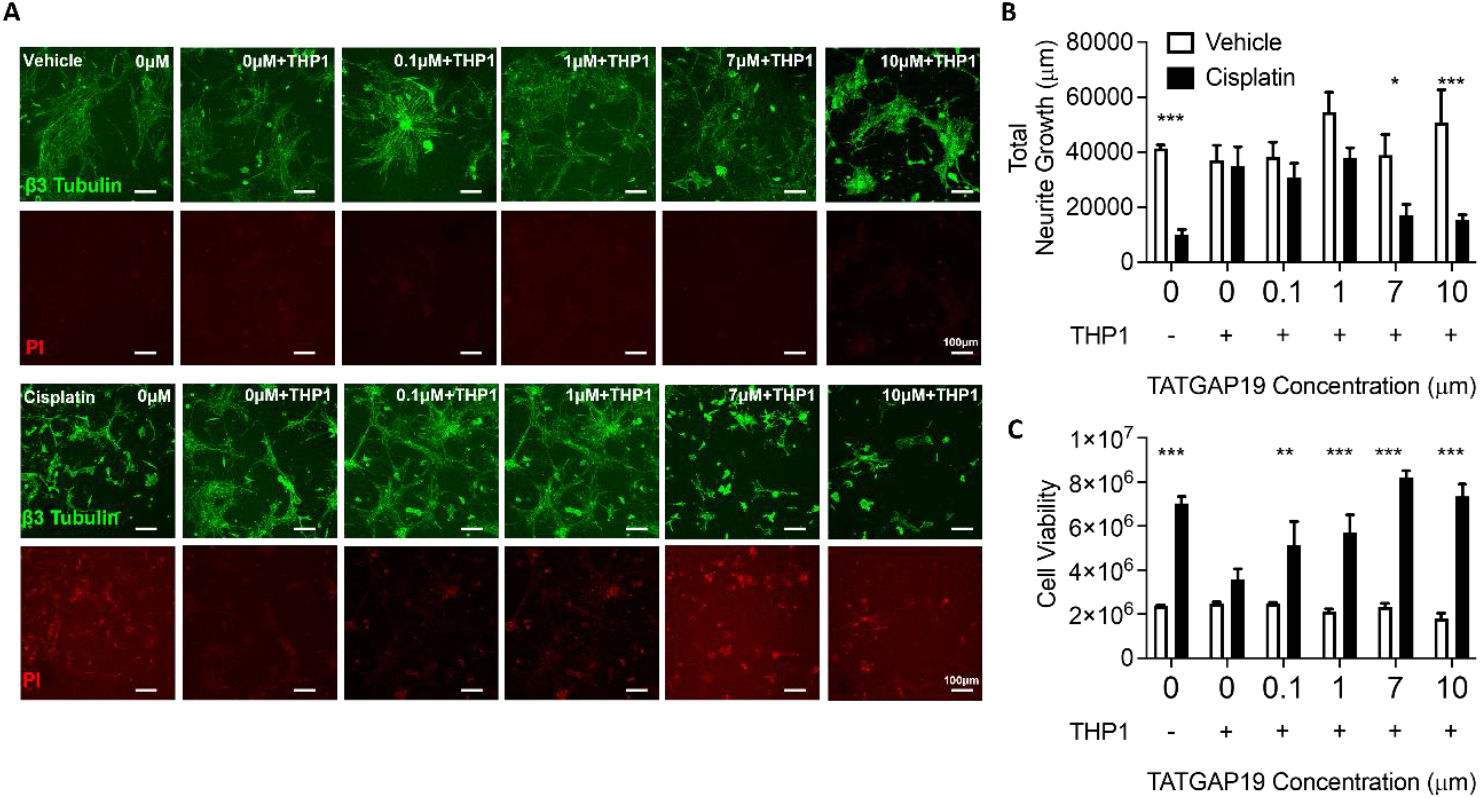
Connexin 43 inhibition prevents mitochondrial transfer from monocytes to DRG neurons recovers neuronal architecture. [A] Representative images of DRG neurons treated with either vehicle or cisplatin and co-cultured with THP-1 cells in response to TATGap19 (Scale bar = 100µm). Neurite growth (green β3 tubulin) and apoptosis (red PI) were determined. [B] An increased neurite length was observed in cisplatin treated DRG neurons following the uptake of monocyte-derived mitochondria, which was prevented by connexin 43 inhibition (**** p<0.0001, one-way ANOVA and a Dunnett’s post-hoc test for multiple comparisons). [C] Decreased cell death (Propidium Iodide) was observed in cisplatin-treated DRG neurons following the uptake of monocyte-derived mitochondria. This was prevented following connexin 43 inhibition. Two-way ANOVA and a Kruskal Wallis test for multiple comparisons. *** P<0.001.

## Discussion

Systemic treatment with cisplatin invokes a general inflammatory response, causing the influx of immune cells into peripheral tissue. The monocytes that infiltrate this tissue release a cascade of pro-inflammatory mediators that influence the functionality of cells within the affected areas [28]. Importantly, neurons are a cell type that are profoundly affected by inflammation induced by cisplatin resulting in neuronal damage leading to peripheral neuropathies[11,23]. Although acute neuropathic pain may be attributed to local mediator release, this is less likely to result in progressively elevated long-term nociception as mediators released are often transitory with traditional analgesics lacking clinical efficacy. This suggests an alternative mechanism driving chronic pain pathology[17]. This has led to a current focus on sensory neuronal mitochondrial dysfunction having a key role in CIPN[30], through dysfunctional oxidative phosphorylation in neuronal mitochondria exposed to cisplatin[25]. This arises as neurons are highly metabolically active with greater ATP demands resulting in a higher density of mitochondrial than normal cells[9]. Therefore, damage to mitochondria following cisplatin treatment will significant impact sensory neuronal functionality and nociception through an oxidative stress driven mechanism. Recent evidence suggests that the drive for macrophages into neuronal tissue such as the dorsal root ganglia is a supportive one, with macrophages donating healthy mitochondria to neuronal cells resulting in an amelioration of pain[29].

This study demonstrates that monocytes transfer mitochondria to cisplatin-damaged sensory neurons to recover neuronal mitochondrial function, with the aim to stabilize neuronal energy balance and alleviate dysfunction that can lead to chronic neuropathic pain. The mechanism that monocytes transfer mitochondria to damaged neurons is through a gap junction mediated mechanism driving a recovery in the architecture of cisplatin-damaged sensory neurons. There is a dynamic balance between mitochondrial fission and fusion that is crucial for maintaining neuronal mitochondrial function and integrity. In the pathophysiology of both inflammatory and neuropathic pain, an essential role of Drp1 in driving mitochondrial fission was uncovered[31]. Here we showed that cisplatin caused an upregulated neuronal mitochondrial fission in cisplatin-damaged sensory neurons producing fragmented mitochondria. This has been associated with impairment in neuronal mitochondrial Ca2+ channels, leading to decreased nociceptor excitability, and the potentiation of chronic neuropathic pain.

We also observed damage to the neuronal mitochondrial network and decreased mitochondrial localization in the neurites of cisplatin-treated DRG neurons in comparison to vehicle treated controls, with mitochondria primarily localized within the cell body. This is a result of the inhibition of microtubule assembly induced by cisplatin, causing the disruption of neuronal mitochondrial trafficking[26]. Additionally, cisplatin exposure resulted in dysmorphic neuronal mitochondria which had a decreased surface area which is consistent with the pathophysiological features of mitochondrial damage and are indicative of a compromised neuronal bioenergetic state.

Cell-cell coupling, and the intercellular exchange of ions, metabolites, and cellular organelles, including mitochondria, critically depend on the involvement of Cx43 in the formation of gap junctions (GJs) and hemichannels (HCs)[7]. A model for the transfer of mitochondria from cells to cells has been proposed to involve the internalization of gap junctions containing Cx43.

Mitochondria present within cellular protrusions that connect to other cells by invaginating gap junctions can become fully enclosed within internalized gap junctions in a receiving cell facilitating mitochondrial transfer. To assess the involvement of Cx43 in mitochondrial transfer between monocytes and cisplatin-damaged neurons, we inhibited the functional properties with a Cx43 inhibitor, Gap-19. This showed a dose-dependent inhibition of mitochondrial transfer from monocytes to cisplatin-damaged sensory neurons confirming Cx43 involvement in the process. These observations provide a promising mechanism for therapeutic target of chronic pain in adult survivors of paediatric cancers.

## Author contribution statement

RPH, MPC, BO, JC performed the experimental work and contributed to the conception or design of the work in addition to acquisition, analysis or interpretation of data for the work. All authors drafted the article or revised it critically for important intellectual content. All authors approved the final version of the manuscript.

## Acknowledgements

This work was supported by the Medical Research Foundation (MRF-RGM-CACP-23-104 to RH and MPC) and Nottingham Trent University. The authors would like to thank Graham Hickman of the Imaging Suite at Nottingham Trent University for their support and assistance in this work.

## Data availability statement

All data is presented in this article and is available by reasonable request from the corresponding author.

## Conflict of interest disclosure

Authors have nothing to disclose.

## Ethics approval statement

This work has not been published or under review for publication elsewhere. All authors have seen and approved the final version of the abstract. All studies involving rodents were approved by local ethical review board at host institutions and in accordance with UK Home Office Legislation.

## Notes

### Competing Interest Statement

The authors have declared no competing interest.

### Summary of Updates

manuscript updated with new introduction, methods and results sections.

